# Fenoxaprop-p-ethyl Susceptibility and Mutation Point Detection of Acetyl-CoA Carboxylase (ACCase) in Different Wild oat (*Avena fatua* L.) Populations from China

**DOI:** 10.1101/549402

**Authors:** Jun-jie Liu, Liuyang Lu, Bai-zhong Zhang, Xi-ling Chen

**Affiliations:** Henan Institute of Science and Technology, Xinxiang 453003, People’s Republic of China

**Keywords:** *Avena fatua*, Acetyl-coa Carboxylase (ACCase), fenoxaprop-p-ethyl resistance, mutation points

## Abstract

To explore resistant mechanism of wild oat to fenoxaprop-p-ethyl, the susceptibility of Acetyl-CoA Carboxylase (ACCase) from 24 wild oat populations to fenoxaprop-p-ethyl, the level of gene expression, and mutation site of ACCase were conducted. *In vitro* ACCase activities were solated and measured by enzyme-linked immunosorbent assay kit (ELISA) assays, the results indicated that the IC_50_ value of the ACCase of the most unsusceptible to fenoxaprop-p-ethyl in the wild oat population from Yexian2017 (W24) was 7206.557-fold compared to that of the ACCase of most susceptible to fenoxaprop-p-ethyl in the wild oat population from Queshan (W11). The differential expression of genes in wild oat treated by the IC_50_ fenoxaprop-p-ethyl concentration (6.9 mg/L) for 24 hours using RNA-seq, digital gene expression (DGE) profling was conducted. We found that 8 unigenes annotated as ACCase genes, 0 up-regulaed expression and 3 down-regulated expression were observed. The down-regulaed expressed *ACCase* was selected for qPCR in the relative susceptible population were significantly more suppressed than the three relative resistant populations. The mutations point of ACCase, Ile-1781-Leu, Trp-1999-Cys, Trp-2027-Cys, Ile-2041-Asn, Asp-2078-Gly, Cys-2088-Arg published were not found in the populations tested by multiple sequence alignment with a model complete ACCase sequence of *Alopecurus myosuroides*. These findings suggest that ACCase plays a critical role in the development of wild oat resistance to fenoxaprop-p-ethyl.

---

Wild oat (*Avena fatua* L.) also named oat grass, belongs to the Gramineae Avena and matures one or two years (Christoffers *et al*. 2002). Wild oat is as malignant weeds that harms wheat, oilseed rape, and other crops in China, and has developed serious resistance to herbicides all over the world (Cavan *et al*. 2001).

At present, wild oat is controlled using chemical herbicides. The herbicide varieties that effectively control wild oat are few. The only inhibitors are metsulfuron-methyl and fenoxaprop-p-ethyl. Fenoxaprop-p-ethyl is a kind of aryloxyphenoxypropionate herbicide (AOPP). Its mode of action is by inhibiting the activity of acetyl-coenzyme A carboxylase (ACCase) in grassy weeds and then blocking normal fatty acid synthesis in plants (Devine & Shukla 2000). As a highly effective herbicide, it has been widely used in wheat fields to prevent grassy weeds, such as wild oat. However, both long-term and single uses of fenoxaprop-p-ethyl in wheat fields have caused resistance to wild oat. In addition, fenoxaprop-p-ethyl only acts on ACCase, so weeds are easily able to develop resistance (Delye 2005). Fenoxaprop-p-ethyl belongs to the high-risk level in the resistant risk classification, in which weeds will typically develop resistance after continuous use for a year (Chen 2018; Pornprom 2006). Recently, wild oat had been difficult to control in wheat fields using only fenoxaprop-p-ethyl. However, the resistance of wild oat to fenoxaprop-p-ethyl has been seldom reported, especially the resistance mechanism.

Carboxylation of acetyl-CoA includes mainly two processes which is involved with carboxylation of biotin and transfer of carboxylation reaction respectively (Konishi *et al*. 1996), the ACCases of plants comprise homogeneity (ACCase I) and heterogeneity (ACCase II), they are both responsible for catalysing acety-CoA to malony-CoA (*Schulte et al.* 1997). Homomeric ACCase localizes in cytosol and it is a pivotal enzyme to the step of fatty acid synthesis reaction. Heteromeric ACCase used to be in cytosol, its production malony-CoA is mainly utilized for elongating fatty acid chain and for synthesizing metabolite associated with flavonoids, resistance-endowing (Shorrosh *et al*. 1994). However, there are two special cases in cellular distribution of the two isoenzymes. The first one is that the chloroplast of rape has not only homomeric ACCase and heteromeric ACCase. The second one is that the ACCases of gramineous plants are all classified as homogeneity whether they present in cytosol or in plastid (Sasaki *et al*. 1997).

Target-site herbicide resistance (TSR) and non-target-site herbicide resistance (NTSR) are the main mechanisms of weed resistance to herbicides. TSR involves a target-site mutation that affects the target enzyme sensitivity and results in much lower or higher expression levels of the target enzyme. Studies of NTSR have mainly included osmosis, reduced herbicide absorption and conduction, metabolic detoxification, shielding effects and buffer functioning to increased herbicide resistance herbicide heightening (Powles *et al*. 1997). ACCase is used as the target of AOPP. Weeds develop resistance to AOPP mainly due to mutations of ACCase; but the overexpression of ACCase in weeds also results in the development of resistance to herbicides (Zhang & Powles 2006). *Sorghum halepense* had developed resistance to enoxaprop-p-ethyl and sethoxydim due to the overexpression of *ACCase* gene (Parker *et al*. 1990; Bradley *et al*. 2015).

Previous studies of the resistance mechanism to ACCase herbicides in resistant Avena spp. populations revealed that ACCase gene mutation can confer the resistance to weed frequently (Maneechote *et al*. 1994, 1997; Seefeldt *et al*. 1996; Shukla *et al*. 1997; Cocker *et al*. 2000). Ile-1781-Leu, Trp-1999-Cys, Trp-2027-Cys, Ile-2041-Asn, Asp-2078-Gly and Cys-2088-Arg are now known mutation sites of ACCase to be associated with resistance to fenoxaprop-p-ethyl in wild oat (Christoffers *et al*. 2002; Liu *et al*. 2007; Cruz-Hipolito *et al*. 2011; Beckie *et al*. 2012). Therefore, the susceptibility of Acetyl-CoA Carboxylase (ACCase) from 24 wild oat populations to fenoxaprop-p-ethyl, digital gene expression (DGE) profling, the level of gene expression, and mutation site of ACCase were conducted in *Avena fatua*. It would provide a theoretical basis for understanding the molecular mechanisms of weed resistance.

## MATERIALS AND METHODS

### Seeds of wild oat

Seeds of wild oat collected from Henan Province were distributed in Xiangxian (Xuchang)=W1, Huojia (Xinxiang) =W2, Huixian (Xinxiang) =W3, Wuzhi (Jiaozuo) =W4, Xiuwu (Jiaozuo) =W5, Xunxian (Hebi)=W6, Fugou (Zhuokou) =W7, Wuyang (Luohe)=W8, Lankao (Kaifeng)=W9, Heshan (Hebi) =W10, Queshan (Zhumadian)=W11, Qixian (Hebi)=W12, Suiyang (Shangqiu)=W13, Zhaoling (Luohe)=W14, Huaiyang (Zhoukou)=W15, Xihua (Jiaozuo)=W16, Dengfeng (Zhengzhou)=W17, Suiping (Zhumadian)=W18, Linying (Luohe)=W19, Sheqi (Nanyang)=W20, Xihua (Zhoukou) =W21, Yanling (Xuchang)=W22, Tangyin (Anyang)=W23, Yexian (Pingdingshan) =W24 in 2017 and Yexian (Pingdingshan) in 2016 = W25.

### Cultivation of seedlings

The greenhouse potting methods of were adopted (Li *et al*. 2010). The seeds of different populations of wild oat were sown into pots with a surface area of 75 cm^2^. The soil surface with the unused herbicide was mixed with a proportion of grass biochar, sifted and cultured in the greenhouse. Rearing conditions were 20°C in the daytime and 15°C at night, 75 ± 5% relative humidity, and a 12:12 h light: dark photoperiod. Seeds of wild oat collected from Henan Province were distributed in 24 populations.

### Chemicals

The test herbicide fenoxaprop-p-ethyl oil formulation (69%) and the active compound of fenoxaprop-p-ethyl (95.8%) were supplied by Noposion (Shenzhen, China). Flavine adenine dinucleotide (FAD), thiamine pyrophosphate (TPP), phenylmethanesulfonyl fluoride (PMSF), sodium pyruvate (C_3_H_3_NaO_3_), alpha naphthol (α-naphthol), Tris (hydroxymethyl) aminomethane hydrochloride (Tris-HCl) and polyvinylpyrrolidone (PVP) were obtained from Sigma-Aldrich (USA). Creatine (C_4_H_9_N_3_O_2_), sodium hydroxide (NaOH), Coomassie brilliant blue–G250, monopotassium phosphate (KH_2_PO_4_), magnesium chloride (MgCl_2_), ethylenediaminetetraacetate acid (EDTA), and bovine serum albumin (BSA) as well as an ACCase-ELISA kit were purchased from Beijing Tongzheng Biological Company (China). HiPure Plant RNA Mini Kit, HiPure Plant DNA Mini Kit and HiPure Gel Pure DNA Mini Kit produced by Magen were used to isolated total RNA, genomic DNA and purify the production of PCR, respectively. Taq-Plus PCR Forest Mix(2x) from NOVA (China) was the reagent for PCR. The kit HiScript®ⅡQRT SsuperMix for qPCR and chamQ™ SYBR® qPCR Master Mix, which are used for cDNA synthesis and qRT-PCR separately, are bought from company Vazyme (Nanjing, China). Peptone, yeast powder and agar powder purchased from AOBOX (Beijing, China) were used make LB culture medium. The Ampicillin purchased from Gentihold (Beijing, China) was applied in *Escherichia coli* transformation experiment as the bacterial inhibitor.

### Instruments and equipment

An ultraviolet and visible spectrophotometer (UV2102PC, Multi Wavelength Fluorescence Spectrophotometer, China), refrigerated centrifuge (5417R, Eppendorf, Germany), and a VMax Microplate Reader (Molecular Devices) with SoftMax Pro 5.4 software were used in the analyses. The primary herbicide sprayer was an atomizer with an auto spray device (Model: ASP-1098, Spray-head: ST110-01, Pressure: 0.2 MPa). The PCR program ran in the Analytik Jena Gradient PCR Instrument (9701 series, Germany) and productions of PCR were validated in the electrophoresis equipment (Beijin Junyi-dongfang Equipment Co.Ltd, Voltage:100v, time: 20 min). The Bio-Rad CFX 96 (USA) was used for qPCR. An constant temperature incubator shaker produced by Jie Rui Er (Jiangsu, China) was used to cultivate *Escherichia coli*.

### In vitro ACCase susceptibility of different wild oat populations to fenoxaprop-p-ethyl

#### Enzyme preparations

Buffer solution: 100 mmol/L Tris-HCl (pH 8.3), 300 mmol/L glycerinum, 5 mmol/L DTT, 2 mmol/L EDTA, 0.5 mmol/L PMSF, and 0.01% (v/v) Triton X-100. First, take 3 g of aerial portions at the 3-leaf stage, tear them in pieces and put them into a prechilled mortar. Grind into a powder in liquid nitrogen, transfer the contents to a 50 mL centrifuge tube, wash the mortar with cold buffer solution in another 50-mL centrifuge tube and place the tube in the refrigerator at 4°C for two hours. Second, primary product to be disposed in two steps. First, put the abovementioned homogenate into a high-speed freezing centrifuge at 4000×g at 4°C for 30 min, discard the sediment, and then, retain liquid supernatant. Second, continue to centrifuge at 20000×g at 4°C for 30 min, discard the sediment, and then, retain the liquid supernatant containing the protein. Transfer the supernatant to a 1.5-mL centrifuge tube to standby for application. Regarding the sample treatment and requirement of the tissue sample, centrifuge the sample for 30 min (4000 r/min). Retain the liquid supernatant for immediate measurement or place it in a −20°C refrigerator for later use. Note that the sample does not contain NaN_3_ because it affects the activity of haptoglobin-related protein (HRP).

#### The ACCase activity of wild oat

The test of the ACCase activity depends on the instruction of the ELISA-ACCase kit. The kit is based on the double antibody-sandwich method for testing the ACCase activity. The depurated ACCase antibody is used to parcel the microplate. Prepare a solid-phase antibody, add the ACCase to the parcel-antibody microplate in the proper order, combine with the ACCase antibody marked by HRP and generate antigen-antibody-enzyme-labelled antibody compounds. After washing the microplate, add the substrate TMB to make it change colours. The TMB turns blue under the catalytic action of the HRP enzyme. The TMB will turn yellow at the end because of the action of the terminating liquid. The depth of the colour is positive correlated with the concentration of ACCase. The spectrophotometric (OD) values are record by a Vmax microplate reader at a wavelength of 450 nm. The protein content was determined by the method of Bradford *et al*. (1976) using bovine serum albumin as a standard. ACCase activity was expressed as U/mg protein/h. The herbicide was applied at a dose rate of 1, 10, 100, 1000, and 10000 mg/L each sample according to preliminary experiments. The inhibition rate of ACCase activity was calculated. The IC_50_ values and the comparative analysis were calculated using a software SPSS 15.0.

#### Fenoxaprop-p-ethyl treatment

A whole-plant assay modified was conducted according to Ryan (1970). Plants thinned and planted in the field (20 plants per pot) at the 3-leaf stage (20 cm plant height), were treated with fenoxaprop-*p*-ethyl by atomizing with an auto spray device (Model: ASP-1098, Spray-head: ST110-01, Pressure: 0.2 MPa). The herbicide was applied at a concentration of 10 g ai/ha (IC_50_) according to the results of a whole plant assay, the amount of spouting liquid was 450 L/ha. Water alone was used as a control. Each treatment was repeated three times. Foliar parts were collected for gene stability analyses at 24 h after treatment and stored at –80°C for RNA extraction.

#### DGE library preparation and sequencing

A total of 10 3-leaf stage, 5 from the control and 5 from fenoxaprop-p-ethyl treated *A. fatua* were collected separately according to the above description, each sample was prepared as 2 biological replicates. RNA extraction was done using TRIzol kit in according to manufacturer instruction, and DNase I (Promega, Madison, WI) was also used to remove genomic DNA from the samples. Approximately 30 μg RNA from each sample was used to construct the DGE libraries, each sample was replicated two times. The mRNA was treated as described in cDNA library construction and enriched by PCR amplification. The library products were then ready for sequencing analysis via Illumina HiseqTM 4000 (Beijing Novogene, Beijing, China) using paired-end technology (150 bp) in a single run. DGE analysis was performed to obtain a global view of wild oat transcriptome differences between the control and fenoxaprop-p-ethyl treated wild oat. To compare differentially expressed genes between the libraries (fenoxaprop-p-ethyl treatment vs. / the control), the level of gene expression was determined by normalizing the number of unambiguous tags in each library to reads per kilobase mapped (RPKM).

#### The effects of fenoxaprop-p-ethyl on the expression of ACCase gene in some populations

Relative expression of ACCase gene in 3 resistant populations and 1 susceptible population (W11) treated by fenoxaprop-p-ethyl at their own IC_50_ concentration in 3-leaf stage were determined. Samples were corrected for qPCR, each treatment was repeated for 3 times. *18S* was selected as the reference gene (Zhang *et al*. 2018), an ACCase sequence from RNA-seq of wild oat was used to design primers by primer 3.0 (http://bioinfo.ut.ee/primer3-0.4.0/). The primers used were listed in Table 1.

**Table 1.**
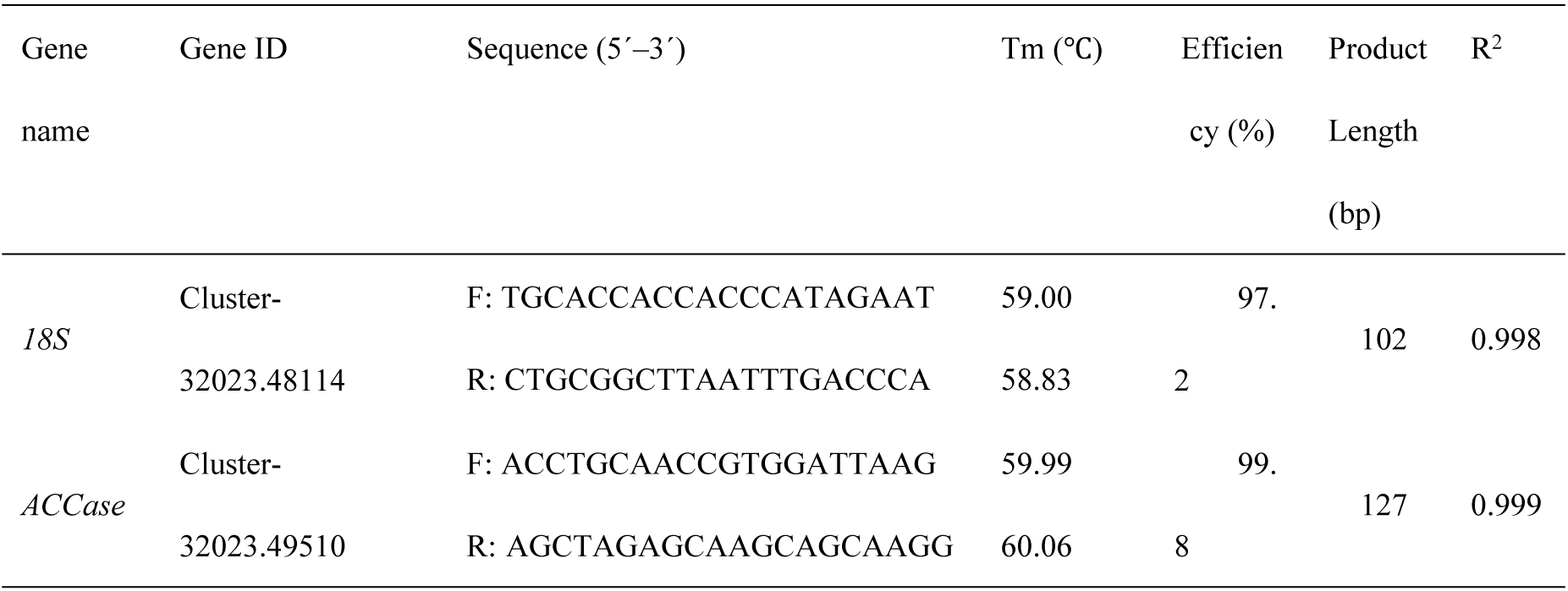
Primer design for qPCR.

#### RNA extraction and cDNA synthesis

To synthesize cDNA, RNA was isolated using TRIzol reagent (Invitrogen, Carlsbad, USA) according to the manufacturer’s instructions. The concentration and quality of RNA was test by NanoDrop 2000 (Thermo Scientific, USA). RNA samples with an A260/A280 ratio ranging from 1.8 to 2.0 and an A260/A230 ratio >2.0 were used for cDNA synthesis. Total RNA (1 µg) was reverse transcribed into First-strand complementary DNA using a PrimeScript RT reagent kit with gDNA Eraser (Takara, Dalian, China) according to the manufacturer’s instructions and stored at –20°C.

#### Quantitative Real-Time PCR (qPCR)

ROX’s Platinum SYBR Green qPCR SuperMix-UDG kit (Invitrogen) was used for qPCR and implementing on an Applied Biosystems 7500 Real-Time PCR system (Applied Biosystems, Foster City, CA). The reactions were performed in a 20 µl volume of a mixture containing 1 µl of cDNA template, 10 µl of SYBR Green qPCR SuperMix-UDG, 0.3 µl of each primer, and 8.7 µl of nuclease-free water. The thermocycling program was as follows: 50°C for 2 min, 95°C for 2 min, and 40 cycles of 95°C for 15 s, and 55°C for 30 s. To acquire a high specificity amplification, a melting curve analysis between 65 to 95°C, was constructed at the end of each PCR run. And it based on a 2-fold dilution series of cDNA (1:5, 1:10, 1:10, 1:10, 1:20, and 1:40). The corresponding qPCR efficiencies (E) were calculated refer to the formula E = 10-1/slope-1 (PFAFFL, 2001; Tellinghuisen 2014; Spiess *et al*. 2015, 2016). Each sample was prepared as 3 biological replicates, and each reaction was analysed with 2 technical replications.

### Mutation point detection

#### DNA extraction, PCR amplification, and cloning

cDNA was isolated from 3-leaf stage issue of 1 relative susceptible populations and 7 relative unsusceptible populations using HiPure Plant DNA Mini Kit. 20 plants was tested in each population selected as mutation point detection. As the complete ACCase sequence is unknown, so PCR primers was designed from *Alopecurus myosuroides* (GeneBank accession No. AJ310767) and the region of PCR product in the wild oat plastidic ACCase carboxyl transferase domain with 5 known and potential ACCase resistance mutation sites, lle-1,781-Leu, Trp-1,999-Cys, Trp-2,027-Cys, lle-2041-Asn, and Asp-2,078-Gly. The PCR was conducted in 25 μL volume that consist of 12.5 μL Taq-Plus PCR Forest Mix(2x) (NOVE, Jiangsu, China), 0.5 μL forward primer: CTGAATGAAGAAGACTATGGTCG, 0.5 μL reverse primer: TCCTCTGACCTGAACTTGATCTC, 1 μL DNA, 10.5 μL dd H_2_O. The PCR was run with the following profiles: 94℃ for 2 min; 25-35 cycles for 30 sec, 60℃ for 30 sec, 72℃ 2 min; 72℃ 5 min. The PCR products was purified from agarose gels using HiPure Gel Pure DNA Mini Kit. The DNA that it had been purified was connected with pGEM-Teasy vector by T4 DNA Ligase as follows: 3 μL PCR product, 5 μL 2 ⅹ T4 Ligase buffer, 1 μL pGEM-Teasy vector, 1 μL Ligase. All of operations above were on the ice and the action was at 4℃ overnight for the maximum number of transformants.

The transformation of ligation product consists mainly of following steps: First, centrifuging the ligation reactions briefly and adding 2 μL of each ligation reaction to a sterile 1.5 mL tube on ice; Second, placing the competent cells in an ice bath until just thawed (5 minutes) and mixing cells by gently flicking the tube; Third, transferring carefully 50 μL cells to the ligation reaction tubes from Step 1, then flicking gently the tubes and incubating on ice for 30 minutes; Fourth, heat-shocking the cells for 45-50 seconds in a water bath at exactly 42℃ without shaking and returning immediately the tubes to ice for 2-3 minutes; Fifth, adding 950 μL room temperature liquid LB medium (without ampicillin) to the ligation reaction transformations and incubating for 1.5 hours at 37℃ with shaking (150-200 rpm); Sixth, plating 100 μL transformation culture onto solid duplicate LB/ampicillin/ IPTG/X-Gal plates; Finally, incubating plates overnight at 37℃.

#### ACCase gene sequencing and alignment

Selecting white colonies and transferring it into liquid LB medium (including ampicillin) continue to incubate for 18 hours at 37℃ with shaking (220 rpm), then it was sequenced by a gene corporation named Beijing Genomic Institute, BGI (Beijing, China). These sequences were analysis and alignment using a software, DNAMAN with a model ACCase sequence (*Alopecurus myosuroides* Gene Bank Accession No. AJ310767) due to the lack of complete wild oat ACCase sequence (Brown 1990)

#### Statistical analysis

Statistical and bioassay analyses were performed using Microsoft Excel (2010). IC_50_ values were calculated by SPSS (15.0). Statistical analysis was performed using one-way analysis of variance and Tukey’s test (*P* < 0·05) with InStat Version 3.0 software (GraphPad Software, San Diego, CA).

## RESULTS

### In vitro ACCase susceptibility of different wild oat populations to fenoxaprop-p-ethyl

The effects of fenoxaprop-p-ethyl on *ACCase* of different wild oat populations is shown in table 2. The results indicate that The most unsusceptible ACCase to fenoxaprop-p-ethyl was found in the wild oat population from Yexian2017 (W24), its IC_50_ value was 8172.236 mg/L, the next unsusceptible ACCase to fenoxaprop-p-ethyl were from Fugou (W7), W21(Xihua), Huixian (W3) and Linying (W19), their IC_50_ values were 7656.177, 5111.930, 3966.196, and 2736.872 mg/L, respectively. The most susceptible ACCase to fenoxaprop-p-ethyl was found in the wild oat population from Queshan (W11), its IC_50_ value was 1.134 mg/L. The IC_50_ of the most unsusceptible ACCase to fenoxaprop-p-ethyl of wild oat population from Yexian2017 (W24), which was 7206.557-fold greater than that of the most susceptible ACCase to fenoxaprop-p-ethyl of the wild oat population from Queshan (W11). In addition, the susceptibility of ACCase to fenoxaprop-p-ethyl of the wild oat from Yexian in 2017 (Pingdingshan) is less than that in 2016; the IC_50_ values were 8172.236 and 1339.554 mg/L, respectively, with an 6.101-fold difference.

**Table 2.** The susceptibility of ACCase to fenoxaprop-p-ethyl on different populations of wild oat.

| oat |  |  |  |  |  |
| --- | --- | --- | --- | --- | --- |
| Populations | SLOPE±S.E | IC<sub>50</sub> (95% Confidence limit)(mg/L) | X<sup>2</sup> | Degree of freedom<br>(Df) | R<sup>2</sup> |
| Xiangxian (W1) | 0.08±0.016 | 115.315 (24.337-447.658) | 7.039 | 13 | 0.971 |
| Huojia (W2) | 0.07±0.023 | 69.285 (43.449-149.531) | 4.050 | 13 | 0.922 |
| Huixian (W3) | 0.05±0.014 | 3966.196(4511.583-5253.136) | 1.573 | 13 | 0.844 |
| Wuzhi (W4) | 0.12±0.038 | 2.725 (1.716-6.723) | 2.889 | 13 | 0.868 |
| Xiuwu (W5) | 0.09±0.034 | 98. 781 (65.103-155.183) | 1.282 | 13 | 0.925 |
| Xunxian (W6) | 0.10±0.040 | 12.065 (7.904-70.572) | 3.527 | 13 | 0.965 |
| Fugou (W7) | 0.07±0.037 | 7656.177 (5962.964- 8554.205) | 6.218 | 13 | 0.932 |
| W8(Wuyang) | 0.13±0.039 | 1.255 (0.986-8.655) | 12.939 | 13 | 0.909 |
| W9(Lankao) | 0.10±0.013 | 1140.105 (1244.703-5456.843) | 7.550 | 13 | 0.878 |
| Heshan (W10) | 0.30±0.018 | 13.805 (5.630-29.448) | 16.850 | 13 | 0.969 |
| Queshan (W11) | 0.17±0.037 | 1.134 (0.525-6.959) | 14.761 | 13 | 0.943 |
| Qixian (W12) | 0.27±0.023 | 36.828 (11.222-83.391) | 11.625 | 13 | 0.827 |
| Suiyang (W13) | 0.20±0.044 | 166.126 (31.72 5-209.475) | 2.466 | 13 | 0.925 |
| Zhaoling (W14) | 0.38±0.032 | 7.140 (2.949-22.873) | 31.076 | 13 | 0.939 |
| Huaiyang (W15) | 0.23±0.065 | 10. 797 (2.668-17.5 65) | 5.102 | 13 | 0.946 |
| Wenxian (W16) | 0.11±0.015 | 560. 584 (229.352-883.867) | 6.978 | 13 | 0.858 |
| Dengfeng (W17) | 0.16±0.037 | 507. 665 (117.969-781.807) | 7.065 | 13 | (0.923) |
| Suiping (W18) | 0.16±0.030 | 130.537 (83.338-476. 601) | 5.700 | 13 | 0.991 |
| Linying (W19) | 0.10±0.024 | 2736.872 (1454.222-2563.884) | 0.819 | 13 | 0.927 |
| Sheqi (W20) | 0.08±0.032 | 133.274 (50.615-186.894) | 20.564 | 13 | 0.937 |
| W21(Xihua) | 0.21±0.064 | 5111.930 (4495.158-5918.343) | 4.998 | 13 | 0.908 |
| Yanling (W22) | 0.13±0.045 | 166.984 (105.604-372.714) | 32.600 | 13 | 0.939 |
| Tangyin (W23) | 0.05±0.013 | 400.835 (186.5252-638.301) | 3.640 | 13 | 0.901 |
| Yexian2017<br>(W24) | 0.06±0.020 | 8172.236 (6454.222-9591.452) | 2.109 | 13 | 0.901 |
| Yexian2016<br>(W25) | 0.08±0.019 | 1339.554 (807.785-1800.296) | 1.792 | 13 | 0.992 |

### In vivo ACCase activity of different wild oat populations

The ACCase activity of different wild oat populations is shown in Fig 1. The ACCase activity of the wild oat population from Suiyang (W13) was the highest at 3.278 U/h/mg protein. The next highest ACCase activity were from Zhaoling (W14), Wenxian (W16), Qixian (W12) and Linying (W19), their ACCase activities were 3.245, 3.234, 3.123 and 3.101 U/h/mg protein, respectively. The ACCase activity of the wild oat population from Heshan (W10) was the lowest, which was 0.664 U/h/mg protein. Among them, the highest ACCase activity of wild oat population from Suiyang (W13) was 4.936-fold that of the lowest population from Heshan (W10). In addition, the ACCase activity of wild oat from Yexian (Pingdingshan) in 2017 was higher than that in 2016, which were 2.492 and 1.356 U/h/mg protein, respectively, a 1.838-fold difference.

**Fig. 1.**
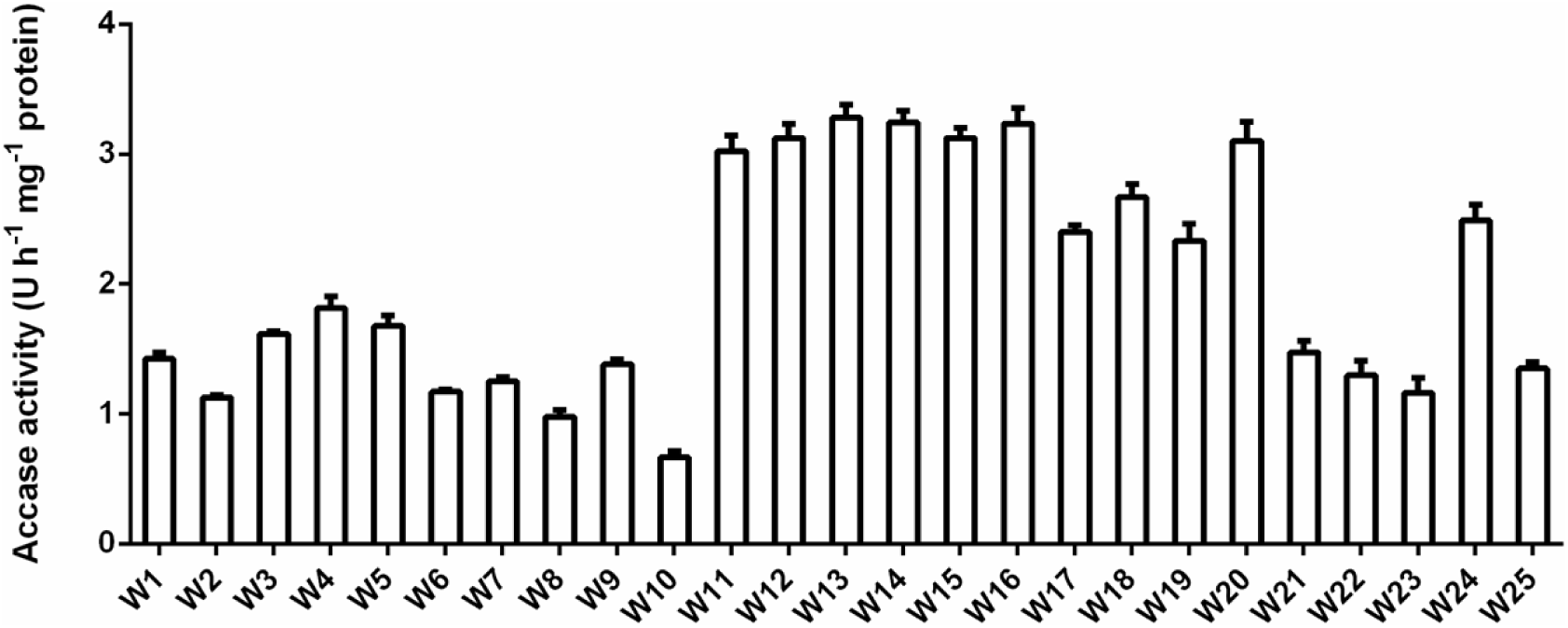
*In vivo* ACCase activity of different wild oat populations.

### Mutation point detection

Eight populations were selected for mutation point detection by multiple sequences alignment and a complete ACCase sequence from *Alopecurus myosuroides* (Gene Bank Accession No. AJ310767), which has high homology with wild oat, was used for the alignment stander. There was no amino acid mutation in six possible sites, it indicated the resistance from these populations in the assay to fenoxaprop-p-ethyl wasn’t caused by the six known amino acid mutations. However, the other sites, Glu-1797-Gly, Thr-1805-Ser, Pro-1829-Leu, Thr-1833-lle, Met-1859-Thr, Asp-1904-Gly, Asn-1913-Asp, Phe-1935-Ser, Gln-2009-Arg, Thr-2092-Ala may be possible amino acid mutations (Fig 2).

**Fig. 2.**
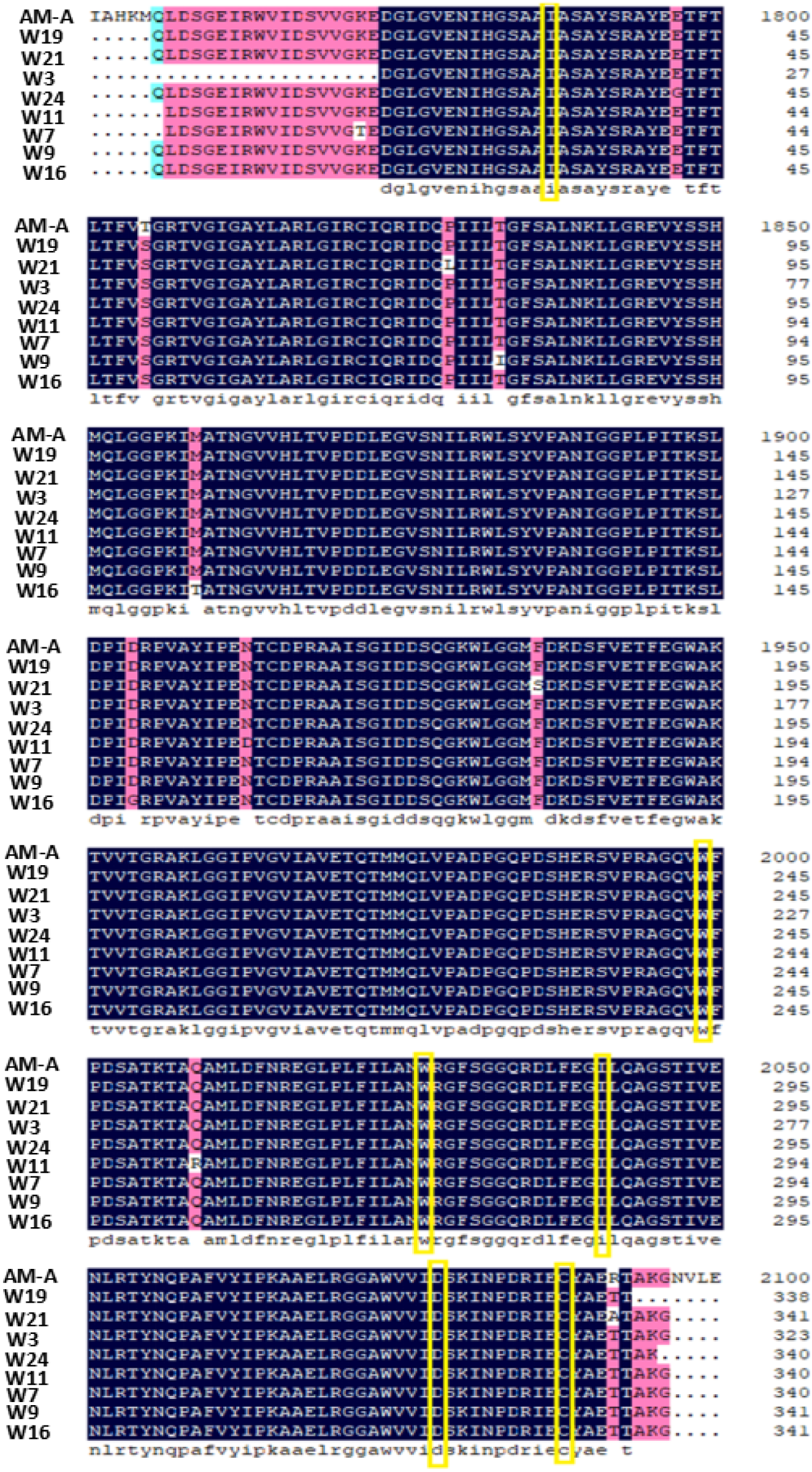
Alignment of the deduced amino acid sequences of ACCase. Yellow frames represent that six possible amino acid mutation site which had been published. AM-A means model ACCase sequence (*Alopecurus myosuroides*). W11 is most susceptive population, the other 7 are relative resistant.

### Differential expressed ACCase unigenes in wild oat suppressed by fenoxaprop-p-thyl

The differential expression was conducted using RNA-seq, digital gene expression (DGE) profling in wild oat treated by the IC_50_ fenoxaprop-p-ethyl concentration (6.9 mg/L) for 24 hours. The results showed that 8 unigenes were annotated as ACCase, 0 up-regulaed expression and 3 down-regulated expression were observed (Table 3).

**Table 3.** Differential expressed ACCase unigenes in wild oat suppressed by fenoxaprop-p-ethyl.

| Gene ID | FTvsCK.log2.<br>(Fold change). | Similarity<br>organisms | Query<br>cover<br>% | E value | Accession |
| --- | --- | --- | --- | --- | --- |
| Cluster-32023.23305 | -1.0028 | *T. aestivum* | 21 | 1e-16 | EU660892.1 |
| Cluster-22064.0 | -0.0028251 | *X. granatum* | 100 | 4e-155 | LC217858.1 |
| Cluster-32023.20930 | -2.0028 | *T. aestivum* | 66 | 2e-84 | EU660895.1 |
| Cluster-32023.63937 | -0.56424 | *T. aestivum* | 63 | 9e-71 | EU660895.1 |
| Cluster-32023.46997 | 0.5603 | *A. tauschii* | 77 | 0.0 | XM_020312452.1 |
| Cluster-32023.62340 | -0.1636 | *A. tauschii* | 96 | 0 | EU660897.1 |
| Cluster-32023.48393 | 0.45485 | *A. myosuroides* | 89 | 0 | AJ632096.1 |
| Cluster-32023.49510 | -1.43894 | *A. myosuroides* | 97 | 0 | AJ310767.1 |

Note: Fold Change (FC) = RPKM of neonicotinoid or sulfoxaflor treated samples/RPKM of untreated samples.
RPKM: Reads per kilo bases per million reads.
FDR ≤ 0.001 and the absolute value of $\text{Log}_2^{\text{FC}} \geq 1$ were used as thresholds to judge the significance of differences in gene expression.

### The effects of fenoxaprop-p-ethyl on the expression of ACCase gene in some populations

The expression of ACCase gene in the 3 resistant populations from Huixian (W3), W21(Xihua), and Yexian2017 (W24), and 1 relative susceptible population from, Queshan (W11) treated by fenoxaprop-p-ethyl were conducted. The results indicated that the expression of ACCase gene from Huixian (W3), Queshan (W11), W21(Xihua), and Yexian2017 (W24), treated by fenoxaprop-p-ethyl was suppressed by 0.24-, 0.81-, 0.14-, and 0.21-fold (Fig 3).

**Fig. 3.**
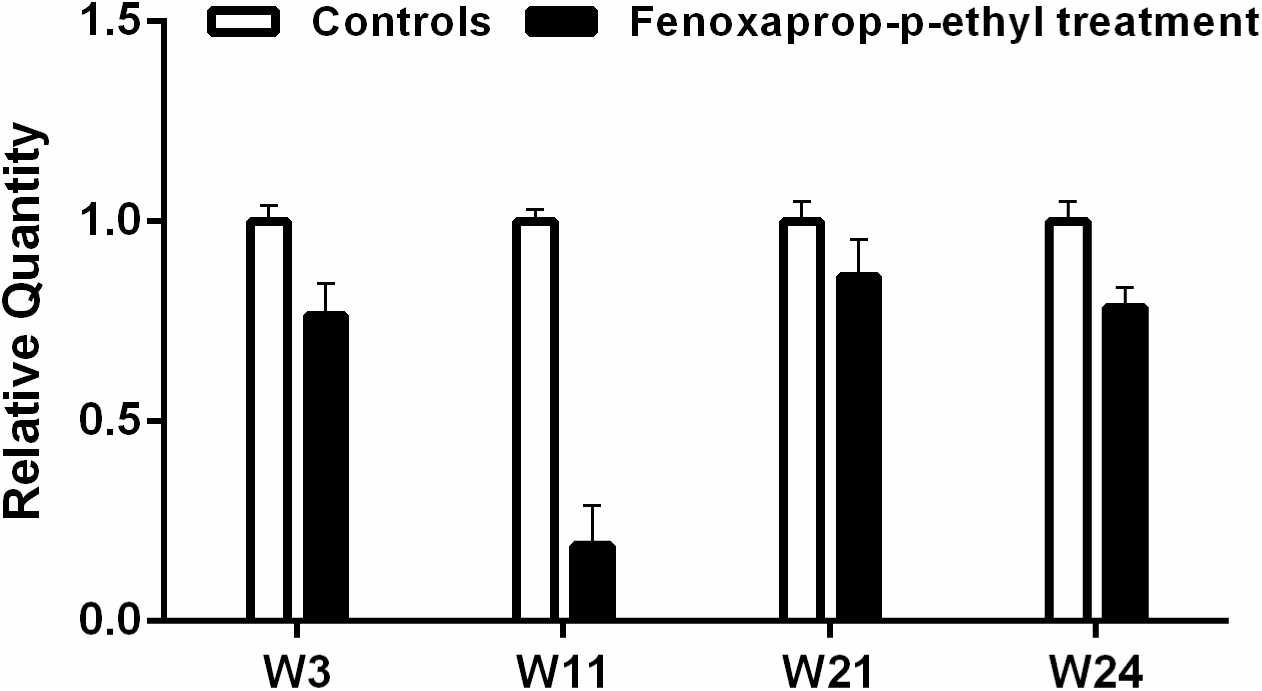
Relative expression of ACCase gene in three resistant populations and a susceptible population (W11). This four typical populations were treated by fenoxaprop-p-ethyl in 3-leaf stage and compared with the control. Date are means ± tandard error (S.E) of three replicates, and the actin gene was used as a reference gene. The relative expression was calculated using 2^−△△Ct^ method base on the value of the control expression, which was ascribed an arbitrary value of 1. W11 is most susceptive population, the other 3 are relative resistant.

## DISCUSSION

Fenoxaprop-p-ethyl is the main ACCase inhibition herbicide used to manage weeds in wheat fields, some species, i.g. *Alopecurus japonicas*, *Beckmannia syzigachne*, and *Avena fatua* have developed different levels of resistance to fenoxaprop-p-ethyl because of its wide usage (Zhang *et al*. 2009; Guo 2011, 2012; Zhang *et al*. 2013; Bi 2013). ACCase is the only target of fenoxaprop-p-ethyl, so the susceptibility of ACCase in plants to fenoxaprop-p-ethyl *in vitro* or *in vivo* could reflect a relationship between the plant biotype and its susceptibility to the herbicide. In this study, susceptibility of ACCase in 24 different populations of wild oat to fenoxaprop-p-ethyl were conducted *in vitro*. The results indicated that the IC_50_ value of the ACCase of the most unsusceptible to fenoxaprop-p-ethyl in the wild oat population from Yexian 2017 (W24) was 7206.557-fold compared to that of the ACCase of most susceptible to fenoxaprop-p-ethyl in the wild oat population from Queshan (W11). This indicated that ACCase of wild oat may play a critical role in the resistance to fenoxaprop-p-ethyl.

Point mutations of ACCase may result in resistance of wild oat to the herbicide, thus six point mutations of ACCase loci published (Christoffers & Berg 2002; Liu *et al*. 2007; Yu *et al*. 2007; Hochberg *et al*. 2009; Kaundun 2010) were determined in the 8 wild oat populations from the 7 relative susceptible and the 1 relative unsusceptible populations to fenoxaprop-p-ethyl, these genetic loci mutations can lead to the different levels of resistance. Amino acid mutations in 1999, 2027, 2041 and 2096 loci of the ACCase of weed species has resulted in resistance to one or more AOPP herbicides but not to cyclohexanedione (CHD) or phenylpyrazoline (PPZ) herbicides, and amino acid mutations in 1781, 2078 and 2088 loci of the ACCase of weed species have led to resistance to all ACCase-inhibiting herbicides (Powles & Yu 2010; Collavo 2011). However, our results indicated that six published resistance-related amino acid mutation sites weren’t found in the populations texted in the eight wild oat populations. However, the other sites, Glu-1797-Gly, Thr-1805-Ser, Pro-1829-Leu, Thr-1833-lle, Met-1859-Thr, Asp-1904-Gly, Asn-1913-Asp, Phe-1935-Ser, Gln-2009-Arg, Thr-2092-Ala may be possible amino acid mutations, so the resistance of wild oat to the herbicide may be caused by other amino sites mutation, ACCase or detoxification enzymes.

The differential expression was conducted using RNA-seq, digital gene expression (DGE) profling in wild oat treated by the IC_50_ fenoxaprop-p-ethyl concentration (6.9 mg/L) for 24 hours. The results showed that 8 unigenes were annotated as ACCase, 0 up-regulaed expression and 3 down-regulated expression were observed. To further clarify the relationship between ACCase and the wild oat resistance to fenoxaprop-p-ethyl, the gene expresssion of ACCase from the 3 resistant populations W21(Xihua), Huixian (W3), Yexian2017 (W24) and the relative susceptible population Queshan (W11) treated by fenoxaprop-p-ethyl for 24 hours were conducted. This indicated that relative expression level of ACCase gene from 3 relative resistant populations and a relative susceptible population had been suppressed by fenoxaprop-p-ethyl, the resistant populations were significantly less suppressed than the susceptible populations.

This proved that the resistance of wild oat to fenoxaprop-p-ethyl is related to the expression level of ACCase gene.

## CONCLUSION

Overexpression of ACCase could play a critical role in the resistance of wild oat to fenoxaprop-p-ethyl. The six published resistance-related amino acid mutation sites weren’t found in the populations texted in the 8 wild oat populations. Resistance mechanisms of wild oat still need further validation, such as other point mutation detection of ACCase, and detoxication enzymes.

## ACKNOWLEDGEMENTS

The authors are highly obliged to the National Key Research and Development Program of China (2017YFD0201700), the Key Science and Technology Program of Henan (Agriculture) (182102110053), the Project of High-level Talent Introduction of Henan Institute of Science and Technology, China (208010616003), the Scientific and Technological Innovation of Henan Institute of Science and Technology, China (208010916005) for the financial support given to the present research work. We thank Z.Y. Huang for reviewing and improving the manuscript.

## ADDITIONAL INFORMATION

The sequence information of Accase unigenes is in the Supplementary S1

## REFERENCES

Bi Y. 2013. Resistance of *Alopecurus japonicus* to fenoxaprop-p-ethyl and mesosulfuron-methyl in winter wheat fields, Shandong Agricultural University.

Bradford M.M.A. 1976. Rapid sensitive method for the quantization of microgram quantities of protein utilizing the principle of protein binding. Analytical Biochemistry 25, 248–254.

Bradley K.W., Wu J., Hatzios K.K. and Esjr H. 2015. The mechanism of resistance to aryloxyphenoxypropionate and cyclohexanedione herbicides in a johnsongrass biotype. Weed Science 49, 477–484.

Brown H.M. 1990. Mode of action, crop selectivity and soil relations of the sulfonylurea herbicides. Pest Management Science 29, 263–281.

Cavan G., Cussans J. and Moss S. 2001. Managing the risks of herbicide resistance in wild oat. Weed Science 49, 236–240.

Chen G., Xu H., Zhang T., Bai C. and Dong L. 2018. Fenoxaprop-P-ethyl resistance conferred by cytochrome P450s and target site mutation in *Alopecurus japonicus*. Pest Management Science 74. doi.org/10.1002/ps.4863.

Christoffers M.J., Berg M.L. and Messersmith C.G. 2002. An isoleucine to leucine mutation in acetyl-CoA carboxylase confers herbicide resistance in wild oat. Génome 45, 1049–1056.

Cocker K.M., Coleman J.O.D., Blair A.M., Clarke J.H. and Moss S.R. 2000. Biochemical mechanisms of cross-resistance to aryloxyphenoxypropionate and cyclohexanedione herbicides in populations of Avena spp. Weed Research 40, 323–334.

Collavo A., Panozzo S., Lucchesi G., Scarabel L. and Sattin M. 2011. Characterisation and management of phalaris paradoxa resistant to ACCase-inhibitors. Crop Protection 30, 293–299.

Delye C. 2005. Weed resistance to acetyl coenzyme A carboxylase inhibitors: an update. Weed Science 53, 728–746.

Devine M.D. and Shukla A. 2000. Altered target sites as a mechanism of herbicide resistance. Crop Protection 19, 881–889.

Guo F. 2011. The resistacne of Japanese foxtail (*Alopecurus japonicus* Steud.) and wild oat (*Avena fatua* L.) to fenoxaprop-p-ethyl and clodinafop-propargyl. Chinese Academy of Agriculatural Sciences.

Guo F. 2012. The sensitivity of different wild oat *Avena fatua* populations to fenoxaprop-P-ethyl and clodinafop-propargyl. Acta Phytophylacica Sinica 39, 87–90.

Hochberg O., Sibony M. and Rubin B. 2009. The response of ACCase-resistant *Phalaris paradoxa* populations involves two different target site mutations. Weed Research 49, 37–46.

Kaundun S.S. 2010. An aspartate to glycine change in the carboxyl transferase domain of acetyl CoA carboxylase and non-target-site mechanism (s) confer resistance to ACCase inhibitor herbicides in a *Lolium multiflorum* population. Pest Management Science 66, 1249–1256.

Konishi T., Shinohara K., Yamada K. and Sasaki Y. 1996. Acetyl-CoA carboxylase in higher plants: most plants other than Gramineae have both the prokaryotic and the eukaryotic forms of this enzyme. Plant Cell Physiology 37, 117–122.

Li R., Zhang J. and Chen G. 2010. Advance of study on identification of weed herbicide resistance. Chinese Agricultural Science Bulletin 26, 289–292.

Liu J., Li P., Lu L., Xie L., Chen X. and Zhang, B. (2018). Selection and evaluation of potential reference genes for gene expression analysis in *Avena fatua*. Plant Protection Science, 55(1), 61–71.

Liu W., Harrison D.K., Chalupska D., Gornicki P., O’Donnell C.C., Adkins S.W., Haselkorn R. and Williams R.R. 2007.Single-site mutations in the carboxyltransferase domain of plastid acetyl-coa carboxylase confer resistance to grass-specific herbicides. Proceedings of the National Academy of Sciences of the United States of America 104, 3627–32.

Liu W., Harrison D.K., Chalupska D., Gornicki P., O’donnell C.C., Adkins S.W., Haselkorn R. and Williams R.R. 2007. Single-site mutations in the carboxyltransferase domain of plastid acetyl-CoA carboxylase confer resistance to grass-specific herbicides. Proceedings of the National Academy of Sciences of the United States of America 104, 3627.

Maneechote C., Holtum J.A.M., Preston C. and Powles S.B. 1994. Resistant Acetyl-CoA Carboxylase is a Mechanism of Herbicide Resistance in a Biotype of *Avena sterilis* ssp. ludoviciana. Plant & Cell Physiology 35, 627–635.

Maneechote C., Preston C. and Powles S.B. 2015. A diclofop - methyl - resistant *Avena sterilis* biotype with a herbicide - resistant acetyl - coenzyme a carboxylase and enhanced metabolism of diclofop - methyl. Pest Management Science 49, 105–114.

Parker W.B., Somers D.A., Wyse D.L., Keith R.A., Burton J.D., Gronwald J.W. and Gengenbach B.G. 1990. Selection and characterization of sethoxydim-tolerant maize tissue cultures. Plant Physiology 92, 1220–1225.

Pfaffl M.W. 2001 A new mathematical model for relative quantification in real-time RT-PCR. Nucleic Acids Research 29, e45.

Pfaffl M.W., Tichopad A., Prgomet C. and Neuvians T.P. 2004. Determination of stable reference genes, differentially regulated target genes and sample integrity: BestKeeper–Excel-based tool using pair-wise correlations. Biotechnology Letters 26, 509–515.

Pornprom T., Mahatamnuchoke P. and Usui K. 2006. The role of altered acetyl-CoA carboxylase in conferring resistance to fenoxaprop-P-ethyl in Chinese sprangletop (*Leptochloa chinensis*, (L.) Nees). Pest Management Science 62, 1109–1115.

Powles S.B., Preston C., Bryan I.B. and Jutsum A.R. 1997. Herbicide resistance: impact and management. Advances in Agronomy 58, 57–93.

Powles S.B. and Yu Q. 2010. Evolution in action: plants resistant to herbicides. Annual Review of Plant Biology 61, 317–347.

Ryan G.F. 1970. Resistance of common groundsel to simazine and atrazine. Weed Science 18, 614– 616.

Schulte W., Töpfer R., Stracke R. and Martini N. 1997. Multi-functional acetyl-CoA carboxylase from Brassica napus is encoded by a multi-gene family: indication for plastidic localization of at least one isoform. Proceedings of the National Academy of Sciences of the United States of America 94, 3465–3470.

Shukla A., Dupont S. and Devine M.D. 1997. Resistance to ACCase-Inhibitor Herbicides in Wild Oat: Evidence for Target Site-Based Resistance in Two Biotypes from Canada. Pesticide Biochemistry & Physiology 57, 147–155.

Seefeldt S.S., Fuerst E.P., Gealy D.R., Shukala A., Irzyk G.P. and Devine M.D. 1996. Mechanisms of resistance to diclofop of two wild oat (*Avena fatua*) biotypes from the Williamette Valley of Oregon. Weed Science 44, 776–781.

Spiess A.N., Deutschmann C., Burdukiewicz M., Himmelreich R., Klat K., Schierack P. and Rödiger S. 2015. Impact of smoothing on parameter estimation in quantitative DNA amplification experiments. Clinical Chemistry 61, 379–388.

Spiess A.N., Rödiger S., Burdukiewicz M., Volksdorf T. and Tellinghuisen J. 2016. System-specific periodicity in quantitative real-time polymerase chain reaction data questions threshold-based quantitation. Scientific Reports 6, 38951.

Tellinghuisen J. and Spiess A.N. 2014. Comparing real-time quantitative polymerase chain reaction analysis methods for precision, linearity, and accuracy of estimating amplification efficiency. Analytical Biochemistry 449, 76–82.

Yu Q., Nelson J.K., Zheng M.Q., Jackson M. and Powles S.B. 2007. Molecular characterisation of resistance to als-inhibiting herbicides in *Hordeum leporinum* biotypes. Pest Management Science 63, 918–927.

Zhang C.X., Ni H.W., Wei S.H., Huang H.J., Liu Y. and Cui H.L. 2009. Current advances in research on herbicide resistance. Scientia Agricultura Sinica 42, 1274–1289.

Zhang C.X., Ni H.W. and Wei S.H. 2013. Herbicide-resistant weeds and their management. Plant Protection 39, 99–102.

Zhang X.Q. and Powles S.B. 2006. The molecular bases for resistance to acetyl co-enzyme A carboxylase (ACCase) inhibiting herbicides in two target-based resistant biotypes of annual ryegrass (*Lolium rigidum*). Planta 223, 550–557.

